# Cabbage stem flea beetle’s (*Psylliodes chrysocephala* L.) susceptibility to pyrethroids and tolerance to thiacloprid in the Czech Republic

**DOI:** 10.1101/584250

**Authors:** Jitka Stará, František Kocourek

## Abstract

The cabbage stem flea beetle (CSFB), *Psylliodes chrysocephala* (Coleoptera: Chrysomelidae), has recently become a major pest species in winter oilseed rape in the Czech Republic. The susceptibility of CSFB populations from two localities to six pyrethroids, two neonicotinoids, one organophosphate and one oxadiazine was evaluated in 2015-2018 in glass vial experiments. The susceptibility of CSFB to thiacloprid and thiamethoxam was evaluated in feeding experiment in 2017 and 2018. High susceptibility of CSFB populations to *lambda*-cyhalothrin, cypermethrin, esfenvalerate, *tau*-fluvalinate, etofenprox, deltamethrin, chlorpyrifos, indoxacarb and acetamiprid was observed in the glass vial experiments. The LC50 and LC90 data obtained for pyrethroids in these experiments in 2015 represent baseline for CSFB resistance monitoring to pyrethroids in the Czech Republic. High tolerance of CSFB to thiacloprid of CSFB was demonstrated in glass vial and the feeding experiment, too.

## Introduction

The cabbage stem flea beetle (CSFB), *Psylliodes chrysocephala* Linnaeus 1758 (Coleoptera: Chrysomelidae) is one of the major pest species of winter oilseed rape in Western Europe [1,2,3]. CSFB has one generation per year and undergoes summer diapause. The period of adult summer diapause is not influenced through constant or changing environmental temperatures or short or long daylight conditions [4]. In autumn, adults migrate into new fields of oilseed rape. Females lay eggs after mating on or close to the plants. After hatching, the larvae feed on leaf bases and terminal leaflets. All the developmental stages can overwinter and females can resume laying eggs in the spring.

Since 2009, the first evidence of CSFB resistance to pyrethroids has been demonstrated as a consequence of the long-term selective pressure of pyrethroids applied for the control of CSFB in oilseed rape [2]. The resistance factor according to the LC50 values for *lambda*-cyhalothrin reached 80 when 28 local populations of CSFB from Germany were examined [1]. Failure to control CSFB has become a major issue in Germany and the United Kingdom [3]. Until now, no data were available about the susceptibility of CSFB to insecticides in the Czech Republic.

Since 2013, three active neonicotinoid ingredients, thiamethoxam, clothianidin and imidacloprid, have been restricted in Europe for seed treatments [5]. Until the start of the restriction, CSFB control in northern Germany and further regions of Western Europe has relied on seed dressings with neonicotinoids or spraying with pyrethroids in autumn [2,6]. Similarly, CSFB in the Czech Republic was effectively controlled until 2013 by seed dressings of winter oilseed rape with neonicotinoids. Using pyrethroids in oilseed rape in autumn was sporadic. In 2014, the first economically important injuries to oilseed rape caused by CSFB were recorded by farmers in the central part of the Czech Republic. Since 2015, CSFB has been controlled in most of the regions of the Czech Republic by spraying with pyrethroids. The present study brings the results of i) evaluation of the susceptibility of local populations of CSFB from the Czech Republic to six pyrethroids, two neonicotinoids, one organophoshate and one oxadiazine in glass vial experiment, ii) evaluation of susceptibility of CSFB to two neonicotinoids in feeding experiments.

## Materials and methods

### Sample collection

The field populations of CSFB originated from winter oilseed rape fields located in Potěhy (49°52′9.695″N, 15°25′12.344″E) and Prague (50°5′15.437″N, 14°17′58.980″E). CSFB adults were collected by sweeping during June in 2015–2018. According to the CSFB population density, 500–1,500 adults per population were collected in particular years and populations. Beetles with some oilseed rape plant material in the net insulators were transported to the laboratory. Prior to initiating the bioassays, samples of the beetles were stored at +5°C. The transported and stored beetles were fed winter oilseed rape leaves. The beetles were tested for susceptibility within 48 h after collection. Only insects in good fitness were used for the bioassays.

### Bioassays

#### Insecticides

Six analytical standards (*lambda*-cyhalothrin, cypermethrin, esfenvalerate, *tau*-fluvalinate, etofenprox and deltamethrin), one organophosphate (chlorpyrifos), one oxadiazine (indoxacarb) and two neonicotinoids (thiacloprid and acetamiprid) were obtained from Sigma Aldrich Co. LLC., Prague, Czech Republic. Four to seven doses of insecticides diluted with acetone were used to generate dose-response curves. The rates recommended in the Czech Republic for the field application were calculated assuming that the highest recommended rate of a commercial product in 400 L of water is used per hectare (Table 1). To generate dose-response curves, the rates of the tested insecticides corresponding to 0.1%-1,000% of the field rates were tested (Table 1). Thiacloprid was tested in 2015-2017 as the analytical standard and in 2017-2018 as the formulated product Biscaya 240 OD. Stock solutions of Biscaya 240 OD were prepared by dissolving 140.4 mg of Biscaya 240 OD formulation (containing 32.4 mg of thiacloprid) in 2 ml of distilled water, and subsequently adjusted to 100 ml with acetone. All further dilutions were made in acetone.

**Table 1.** The recommended field rates and tested rates of insecticides used to evaluate CSFB susceptibility from the Prague and Potěhy localities in 2015-2018.

| Tested insecticides | Field rates ($\mu\text{g}/\text{cm}^2$) | Tested rates ($\mu\text{g}/\text{cm}^2$) |
| --- | --- | --- |
| cypermethrin | 0.25 | 0.0025, 0.025, 0.05, 0.25 |
| deltamethrin | 0.075 | 0.000075, 0.00075, 0.0075, 0.015, 0.075, 0.75 |
| esfenvalerate | 0.075 | 0.0075, 0.015, 0.075, 0.75 |
| etofenprox | 0.6 | 0.06, 0.12, 0.60, 6.00 |
| *lambda*-cyhalothrin | 0.075 | 0.00075, 0.00075, 0.0075, 0.015, 0.075, 0.75 |
| *tau*-fluvalinate | 0.48 | 0.00048, 0.0048, 0.0192, 0.048, 0.096, 0.48, 4.80 |
| chlorpyrifos | 3.00 | 0.03, 0.0752, 0.15, 0.30, 0.60, 3.00 |
| indoxacarb | 0.255 | 0.024, 0.048, 0.24, 2.40 |
| acetamiprid | 0.20 | 0.002, 0.02, 0.04, 0.20 |
| thiacloprid | 0.72 | 0.072, 0.144, 0.72, 1.44, 7.20 |

#### Glass vial experiment

IRAC adult vial tests [7] no. 31 (pyrethroids), no. 21 (neonicotinoids), no. 25 (organophosphates) and no. 27 (oxadiazines) were used to conduct the bioassay. The glass vials (P-lab, CZ, internal surface 32.4 cm^2^) were filled with 520 μl of insecticide and rotated on a hotdog roller at room temperature until the acetone was completely evaporated. Subsequently, the beetles were transferred into each vial. Three replicates were used, and 10 beetles were used for each insecticide dose in each replication. The control vials were treated with pure acetone. The vials were incubated under controlled conditions at 20°C and 60% relative humidity with a 16:8 h light:dark photoperiod. The number of severely affected beetles (i.e., dead and moribund) was counted after 48 h.

#### Feeding experiment

Oilseed rape of the Exssence variety was sown at the rate of 400,000 seeds per ha with a plot seeder Wintersteiger (Wintersteiger Sägen GmbH, Arnstadt, Germany) on small plots (0.2 ha) in the field in Prague locality with three variants: (1) without seed dressing, (2) seed dressing with Sonido (thiacloprid), and (3) seed dressing with Cruiser (thiamethoxam). The plots were left without treatment with herbicides or insecticides. The conditions of the experiment permitted the natural infestation by CSFB adults. In 2017, the plants from the experiment were collected in two periods, 24 days and 35 days after sowing of the oilseed rape seeds. In 2018, the plants were collected 29 days after sowing of the oilseed rape seeds. On each plant collection day, the plants were weighed on a Kern electronic scale EMB200-2 (Kern & Sohn, Balingen, Germany) and the injury caused by CSFB adults to the plants in percentage of leaf loss was evaluated. From 10 to 15 plants in three repetitions from each variant were analysed for their content of thiacloprid and thiamethoxam residues.

### Analysis of residues of neonicotinoids in oilseed rape plants

The residues of thiacloprid and thiamethoxam as active substances of the seed dressings Sonido and Cruiser, respectively, were analysed in samples of oilseed rape plants by the QuEChERS method followed by LC–MS/MS. From each variant and repetition, a composite sample of 2.50 g of plants was used for the analyses. The analyses were carried out in the accredited laboratory at the University of Chemistry and Technology in Prague as a service.

### Data analysis

The bioassay data were analysed using a probit analysis via a dose-effect analysis in XLSTAT 2015 (Addinsoft USA, New York, NY, USA). The doses of insecticides were ln-transformed. The lethal concentrations (LC50s and LC90s) were fitted with 95% confidence limits. The LC50 and LC90 values in particular bioassays were considered significantly different when the respective 95% confidence limits did not overlap.

The mortality data of the CSFB after application of thiacloprid in particular years and localities in the glass vial experiments and the injury of plants, weight of the oilseed rape plants and content of active substances of pesticides in the seed-dressing experiments were compared using a one-way analysis of variance (ANOVA) after testing their normality by the Shapiro-Wilk test using the XLSTAT 2015 statistical software package (Addinsoft Inc., New York, USA). The means were separated using the Tukey HSD analysis.

## Results

### Susceptibility to pyrethroids in the glass vial experiment

The susceptibility of CSFB populations from Prague and Potěhy localities in 2015 – 2018 to 6 active substances of pyrethroids expressed as LC50s and LC90s are given in Tables 2a,b and 3a,b, respectively. The LC50 and LC90 values demonstrated that both tested local CSFB populations were highly susceptible to the pyrethroids cypermethrin, deltamethrin, esfenvalerate, etofenprox, *lambda*-cyhalothrin and *tau*-fluvalinate. The LC50s for the pyrethroids evaluated after 48 h ranged from 0.0002 μg/cm^2^ (deltamethrin, Prague 2016) to 0.13 μg/cm^2^ (*tau*-fluvalinate, Prague 2018). The LC90s ranged from 0.0009 μg/cm^2^ (deltamethrin, Prague 2016) to 0.56 μg/cm^2^ (*tau*-fluvalinate, Prague 2018). The LC50 values for *lambda*-cyhalothrin didn’t differ between years 2015-2018 in CSFB populations from both the localities (Tables 2a and 2b). The LC90 values for *lambda*-cyhalothrin for CSFB from Prague locality was significantly higher in 2017 and 2018 than in 2015, indicating a slight decrease in CSFB susceptibility to *lambda*-cyhalothrin (Table 3a). Similarly, the LC90 value for *lambda*-cyhalothrin for CSFB from Potěhy locality was significantly higher in 2018 than in 2015 (Table 3b). The LC50 and LC90 values for *tau*-fluvalinate for CSFB from Potěhy in 2018 was significantly higher than the values in 2015 (Tables 2b and 3b). Similarly, the LC50 and LC90 values for *tau*-fluvalinate in Prague were higher that the values in 2015 (Tables 2a and 3a). However, the high variability in the data for *tau*-fluvalinate for CSFB from Prague locality in 2015 did not allow the calculation of the confidence limits for the LC50 and LC90 values and comparisons of the CSFB susceptibility between 2015 and following years. The LC50 and LC90 values for *tau*-fluvalinate in 2017 and 2018 in Prague locality were not significantly different (Tables 2a and 3a).

**Table 2a.** Lethal concentrations LC50 (μg/cm^2^) of active substances of insecticides and confidence limits (CL) calculated for populations of CSFB collected in Prague locality in 2015-2018. Values with different numbers for the same letters in the same line are significantly different (the CLs overlap). nd – no CL defined. x – no data available.

| Prague |  |  |  |  |
| --- | --- | --- | --- | --- |
| Active substance | 2015 | 2016 | 2017 | 2018 |
| *lambda*-cyhalothrin | 0.0003 (0.0002/0.0005)<sup>a</sup> | x | 0.001 (0.0004/0.002)<sup>a</sup> | 0.001 (0.0005/0.003)<sup>a</sup> |
| *tau*-fluvalinate | 0.008 (x) | x | 0.09 (0.06/0.13)<sup>a</sup> | 0.13 (0.09/0.19)<sup>a</sup> |
| deltamethrin | 0.0006 (0.0004/0.001)<sup>b</sup> | 0.0002 (0.0001/0.0003)<sup>a</sup> | x | x |
| cypermethrin | 0.006 (0.004/0.009) | x | x | x |
| esfenvalerate | 0.007 (x) | x | x | x |
| etofenprox | 0.10 (0.08/0.14) | x | x | x |
| chlorpyrifos | 0.67 (0.55/0.98)<sup>b</sup> | 0.01 (0.01/0.02)<sup>a</sup> | 0.03 (x) | 0.13 (x) |
| indoxacarb | 0.03 (0.01/0.06)<sup>a</sup> | 0.02 (0.01/0.03)<sup>a</sup> | x | x |
| acetamiprid | 0.03 (0.03/0.04)<sup>b</sup> | 0.002 (0.005/0.003)<sup>a</sup> | x | x |
| thiacloprid | 0.63 (0.32/7.54)<sup>a</sup> | 7.92 (2.78/89.7)<sup>ab</sup> | 139 (67.8/534)<sup>b</sup> | x |
| Biscaya | x | x | 0.72 (0.51/1.008)<sup>a</sup> | 1.48 (0.54/33.7)<sup>a</sup> |

**Table 2b.** Lethal concentrations LC50 (μg/cm^2^) of active substances of insecticides and confidence limits (CL) calculated for populations of CSFB collected in Potěhy locality in 2015-2018. Values with different numbers for the same letters in the same line are significantly different (the CLs overlap). nd – no CL defined. x – no data available.

| Potěhy |  |  |  |  |
| --- | --- | --- | --- | --- |
| Active substance | 2015 | 2016 | 2017 | 2018 |
| *lambda*-cyhalothrin | 0.002 (0.001/0.003)<sup>a</sup> | x | x | 0.004 (0.001/0.007)<sup>a</sup> |
| *tau*-fluvalinate | 0.02 (0.01/0.03)<sup>a</sup> | x | x | 0.08 (0.05/0.12)<sup>b</sup> |
| deltamethrin | 0.002 (0.002/0.003) | x | x | x |
| cypermethrin | 0.02 (0.02/0.03) | x | x | x |
| esfenvalerate | 0.02 (0.01/0.02) | x | x | x |
| etofenprox | 0.08 (0.07/0.10) | x | x | x |
| chlorpyrifos | 0.42 (x) | x | x | x |
| indoxacarb | 0.03 (0.02/0.04) | x | x | x |
| acetamiprid | 0.02 (0.02/0.03) | x | x | x |
| thiacloprid | 8.10 (2.60/152)<sup>a</sup> | 14.1 (3.79/686)<sup>a</sup> | 67.8 (42.9/127)<sup>a</sup> | x |
| Biscaya | x | x | 0.67 (0.50/0.89) | 3.47 (nd) |

**Table 3a.** Lethal concentrations LC90 (μg/cm^2^) of active substances of insecticides and confidence limits (CL) calculated for populations of CSFB collected in Prague locality in 2015-2017. Values with different numbers for the same letters in the same line are significantly different (the CLs overlap). nd – no CL defined. x – no data available.

| Prague |  |  |  |  |
| --- | --- | --- | --- | --- |
| Active substance | 2015 | 2016 | 2017 | 2018 |
| *lambda*-cyhalothrin | 0.001 (0.0008/0.003)<sup>a</sup> | x | 0.009 (0.004/0.03)<sup>b</sup> | 0.01 (0.004/0.03)<sup>b</sup> |
| *tau*-fluvalinate | 0.02 (x) | x | 0.49 (0.28/1.36)<sup>a</sup> | 0.56 (0.33/1.50)<sup>a</sup> |
| deltamethrin | 0.003 (0.002/0.007)<sup>a</sup> | 0.0009 (0.0005/0.002)<sup>a</sup> | x | x |
| cypermethrin | 0.02 (0.01/0.04) | x | x | x |
| esfenvalerate | 0.008 (x) | x | x | x |
| etofenprox | 0.24 (0.17/0.58) | x | x | x |
| chlorpyrifos | 1.29 (0.92/3.33)<sup>b</sup> | 0.03 (0.03/0.04)<sup>a</sup> | 0.04 (x) | 0.19 (x) |
| indoxacarb | 0.53 (0.26/2.15)<sup>b</sup> | 0.08 (0.06/0.17)<sup>a</sup> | x | x |
| acetamiprid | 0.05 (0.04/0.08)<sup>a</sup> | 0.02 (0.01/0.04)<sup>a</sup> | x | x |
| thiacloprid | 13.0 (2.33/nd)<sup>a</sup> | 508 (55.5/nd)<sup>a</sup> | 2,060 (536/nd)<sup>a</sup> | x |
| Biscaya |  |  | 3.20 (2.10/6.25)<sup>a</sup> | 205 (14.7/nd)<sup>b</sup> |

**Table 3b.** Lethal concentrations LC90 (μg/cm^2^) of active substances of insecticides and confidence limits (CL) calculated for populations of CSFB collected in Potěhy locality in 2015-2017. Values with different numbers for the same letters in the same line are significantly different (the CLs overlap). nd – no CL defined. x – no data available.

| Potěhy |  |  |  |  |
| --- | --- | --- | --- | --- |
| Active substance | 2015 | 2016 | 2017 | 2018 |
| *lambda*-cyhalothrin | 0.006 (0.004/0.01)<sup>a</sup> | x | x | 0.04 (0.02/0.11)<sup>b</sup> |
| *tau*-fluvalinate | 0.02 (0.01/0.03)<sup>a</sup> | x | x | 0.43 (0.25/1.16)<sup>b</sup> |
| deltamethrin | 0.01 (0.006/0.02) | x | x | x |
| cypermethrin | 0.07 (0.05/0.13) | x | x | x |
| esfenvalerate | 0.03 (0.02/0.06) | x | x | x |
| etofenprox | 0.16 (0.12/0.32) | x | x | x |
| chlorpyrifos | 0.46 (x) | x | x | x |
| indoxacarb | 0.13 (0.09/0.29) | x | x | x |
| acetamiprid | 0.03 (0.03/0.05) | x | x | x |
| thiacloprid | 822 (66.0/nd)<sup>a</sup> | 1,564 (93.0/nd)<sup>a</sup> | 418 (200/1,605)<sup>a</sup> | x |
| Biscaya |  |  | 2.06 (1.45/3.68) | nd |

The mortalities of CSFB from Prague and Potěhy localities after application of deltamethrin, cypermethrin, esfenvalerate, etofenprox, *lambda*-cyhalothrin and *tau*-fluvalinate in concentrations corresponding to 100% of the recommended field concentration were 100% in 2015 (Tables 4a and 4b). In 2017 and 2018, the mortality of CSFB from Prague locality after application of *tau*-fluvalinate in concentration corresponding to 100% of the recommended field concentration decreased to 86.7% and 79.2%, respectively (Table 4a). However, the difference in CSFB mortalities from Prague locality between years was not statistically significant. Similarly, the mortality of CSFB from Potěhy decreased from 2015 to 2018 to 86.9% and the difference in mortalities was statistically significant (Table 4b). No decrease of CSFB mortality in 2017 or 2018 was observed after application of *lambda-* cyhalothrin in both the localities (Tables 4a and 4b).

**Table 4a.**
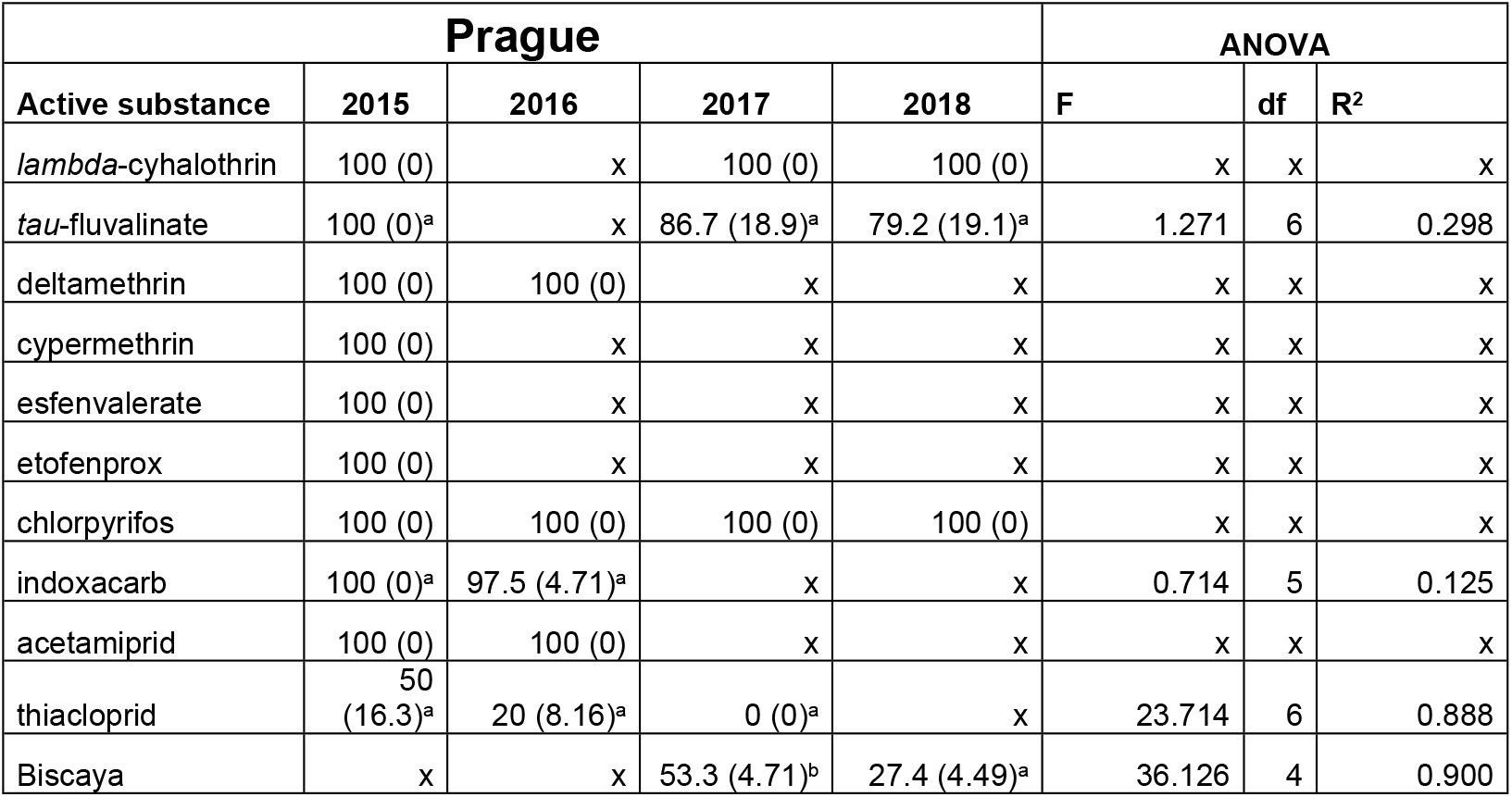
Mean mortality (%) of CSFB from Prague locality at 100% of the recommended field rate of insecticides. Standard deviations are given in brackets. Values with different letters within a row are significantly different. x – no data available.

| Prague |  |  |  |  | ANOVA |  |  |
| --- | --- | --- | --- | --- | --- | --- | --- |
| Active substance | 2015 | 2016 | 2017 | 2018 | F | df | R<sup>2</sup> |
| *lambda*-cyhalothrin | 100 (0) | x | 100 (0) | 100 (0) | x | x | x |
| *tau*-fluvalinate | 100 (0)<sup>a</sup> | x | 86.7 (18.9)<sup>a</sup> | 79.2 (19.1)<sup>a</sup> | 1.271 | 6 | 0.298 |
| deltamethrin | 100 (0) | 100 (0) | x | x | x | x | x |
| cypermethrin | 100 (0) | x | x | x | x | x | x |
| esfenvalerate | 100 (0) | x | x | x | x | x | x |
| etofenprox | 100 (0) | x | x | x | x | x | x |
| chlorpyrifos | 100 (0) | 100 (0) | 100 (0) | 100 (0) | x | x | x |
| indoxacarb | 100 (0)<sup>a</sup> | 97.5 (4.71)<sup>a</sup> | x | x | 0.714 | 5 | 0.125 |
| acetamiprid | 100 (0) | 100 (0) | x | x | x | x | x |
| thiacloprid | 50<br>(16.3)<sup>a</sup> | 20 (8.16)<sup>a</sup> | 0 (0)<sup>a</sup> | x | 23.714 | 6 | 0.888 |
| Biscaya | x | x | 53.3 (4.71)<sup>b</sup> | 27.4 (4.49)<sup>a</sup> | 36.126 | 4 | 0.900 |

**Table 4b.** Mean mortality (%) of CSFB from Potěhy locality at 100% of the recommended field rate of insecticides. Standard deviations are given in brackets. Values with different letters within a row are significantly different. x – no data available.

| Potěhy |  |  |  |  | ANOVA |  |  |
| --- | --- | --- | --- | --- | --- | --- | --- |
| Active substance | 2015 | 2016 | 2017 | 2018 | F | df | R<sup>2</sup> |
| *lambda*-cyhalothrin | 100 (0) | x | x | 100 (0) | x | x | x |
| *tau*-fluvalinate | 100 (0)<sup>a</sup> | x | x | 86.9 (4.44)<sup>b</sup> | 98.885 | 4 | 0.961 |
| deltamethrin | 100 (0) | x | x | x | x | x | x |
| cypermethrin | 100 (0) | x | x | x | x | x | x |
| esfenvalerate | 100 (0) | x | x | x | x | x | x |
| etofenprox | 100 (0) | x | x | x | x | x | x |
| chlorpyrifos | 100 (0) | x | x | x | x | x | x |
| indoxacarb | 100 (0) | x | x | x | x | x | x |
| acetamiprid | 100 (0) | x | x | x | x | x | x |
| thiacloprid | 43.3 (9.43)<sup>a</sup> | 26.7 (23.5)<sup>a</sup> | 0 (0)<sup>a</sup> | x | 5.233 | 5 | 0.677 |
| Biscaya | x | x | 33.3 (5.77)<sup>a</sup> | 16.3 (15.5)<sup>a</sup> | 1.797 | 4 | 0.310 |

The increased LC50 and LC90 values for *tau*-fluvalinate in 2017 and 2018 together with the decreased mortality after application of *tau*-fluvalinate in 2017 and 2018 indicates a shift in CSFB susceptibility to *tau*-fluvalinate after three years of CSFB control by spraying *tau*-fluvalinate in the spring period onto oilseed rape.

### Susceptibility to organophosphates in the glass vial experiment

Sufficient susceptibility of CSFB from two localities to chlorpyrifos was demonstrated in 2015-2018. The LC50 values ranged from 0.01 μg/cm^2^ (Prague 2016) to 0.67 μg/cm^2^ (Prague 2015) and the LC90 values ranged from 0.03 μg/cm^2^ (Prague 2016) to 1.29 μg/cm^2^ (Prague 2015). The high variability in the data from Potěhy in 2015 and Prague in 2017 and 2018 did not allow calculation of the confidence limits for the LC50 and LC90 values (Tables 2a,b and 3a,b).

The CSFB mortality after application of chlorpyrifos in concentration corresponding to 100% of the recommended field concentration was 100% in both the localities (Tables 4a and 4b). Data obtained for chlorpyrifos in 2015-2018 demonstrate stable high CSFB susceptibility from two localities to organophosphates.

### Susceptibility to oxadiazine in the glass vial experiment

The LC50 and LC90 values for CSFB from two localities in 2015-2016 showed a high susceptibility to indoxacarb (Tables 2a,b and 3a,b). The LC50s for indoxacarb ranged from 0.02 μg/cm^2^ (Prague 2016) to 0.03 μg/cm^2^ (Prague 2015, Potěhy 2015) and did not differ significantly between populations. The LC_90_s ranged from 0.08 μg/cm^2^ (Prague 2016) to 0.53 μg/cm^2^ (Prague 2015). The CSFB mortality after application of indoxacarb in concentrations corresponding to 100% of the field concentration was 100% in 2015 in both the Prague and Potěhy localities. In 2016, the mortality of CSFB from Prague decreased to 97.5% (Table 4a). However, the difference in mortalities between 2015 and 2016 was not significantly different.

### Susceptibility to neonicotinoids in the glass vial experiment

The LC50 and LC90 values indicate that the CSFB populations from two localities were susceptible to acetamiprid. The LC50s ranged from 0.002 μg/cm^2^ (Prague 2016) to 0.03 μg/cm^2^ (Prague 2015) (Tables 2a and 2b). The LC90s ranged from 0.02 μg/cm^2^ (Prague 2016) to 0.05 μg/cm^2^ (Prague 2015) (Tables 3a and 3b). The CSFB mortalities after application of acetamiprid in concentrations corresponding to 100% of the recommended field concentration was 100% in both the Prague and Potěhy localities (Tables 4a and 4b).

In contrast to acetamiprid, the susceptibility of CSFB populations from both the localities to thiacloprid tested as analytical standard was insufficient. The LC50 values for thiacloprid ranged from 0.63 μg/cm^2^ (Prague 2015) to 139 μg/cm^2^ (Prague 2017) (Tables 2a and 2b). The LC90s for thiacloprid ranged from 13.0 μg/cm^2^ (Prague 2015) to 2,060 μg/cm^2^ (Prague 2017) (Tables 3a and 3b). In both localities, comparison of the LC50 values between the years showed a decreasing susceptibility of CSFB to thiacloprid from 2015 to 2017 (Tables 2a and 2b). The LC50 and LC90 values obtained for thiacloprid as a formulated product Biscaya 240 OD were much lower that the values for analytical standard of thiacloprid (Tables 2a and 2b).

The CSFB mortality after application of analytical standard of thiacloprid in concentrations corresponding to 100% of the recommended field concentration decreased from 50.0% in Prague locality in 2015 to 0% in both the localities in 2017 (Tables 4a and 4b). However, the difference was not statistically significant. The mortality data obtained for formulated product Biscaya 240 OD showed decreasing mortalities of CSFB from 2017 to 2018 in both the localities.

### Susceptibility of CSFB to neonicotinoids in the feeding experiment

The content of pesticide residues differed between the treatment variants in both plant collection periods in 2017. In the first period (24 days from seeding), the content of thiamethoxam in plants was 17 times higher than the content of thiacloprid. In the second treatment period (35 days from seeding), the content of thiamethoxam in plants was 5 times higher than the content of thiacloprid (Table 5). In 2018, 29 days from seeding, the content of thiamethoxam in plants was 2 times higher than the content of thiacloprid. The average weight of the plants was significantly higher in the thiamethoxam variant than in the thiacloprid and control variants in the first plant collection period in 2017. In the second plant collection period in 2017 and in 2018, the average weight of plants did not differ significantly between the variants thiacloprid and thiamethoxam. The average injury to plants in the thiacloprid variant did not differ significantly from the control variant without treatment and was significantly higher than in the thiamethoxam variant in both years and all periods of plant collection (Table 5).

**Table 5.** Content of pesticide residues in oilseed rape plants (values ± standard deviation), average weight of plants before analysis and average injury to plants caused by CSFB. Values with different letters within a column in each experiment are significantly different. x – no data available.

| year | days from seeding | variant | average content of pesticide residues (mg/kg) | average weigh of plant (g) | average injury of plant (%) |
| --- | --- | --- | --- | --- | --- |
| 2017 | 24 | control | x | 0.34 ±0.04<sup>a</sup> | 34.7 ±3.27<sup>a</sup> |
|  |  | thiaclopid | 185 ±96.6<sup>a</sup> | 0.28 ±0.02<sup>a</sup> | 46.3 ±10.6<sup>a</sup> |
|  |  | thiamethoxam | 3,213 ±122<sup>b</sup> | 0.57 ±0.03<sup>b</sup> | 0.97 ±0.91<sup>b</sup> |
| 2017 | 35 | control | x | 0.83 ±0.11<sup>a</sup> | 74.7 ±12.1<sup>a</sup> |
|  |  | thiaclopid | 43.7 ±12.2<sup>a</sup> | 0.91 ±0.12<sup>a</sup> | 86.8 ±4.15<sup>a</sup> |
|  |  | thiamethoxam | 208 ±47.8<sup>b</sup> | 1.09 ±0.18<sup>a</sup> | 9.08 ±3.49<sup>b</sup> |
| 2018 | 29 | control | x | 0.84 ±0.32<sup>a</sup> | 37.0 ±21.6<sup>a</sup> |
|  |  | thiaclopid | 44.8 ±0 | 1.80 ±0.90<sup>b</sup> | 37.0 ±16.4<sup>a</sup> |
|  |  | thiamethoxam | 101 ±0 | 1.69 ±0.39<sup>b</sup> | 15.0 ±9.72<sup>b</sup> |

## Discussion

### Susceptibility to pyrethroids in the glass vial experiment

The present study showed good efficacy of the pyrethroids *lambda*-cyhalothrin, deltamethrin, cypermethrin, *tau*-fluvalinate, esfenvalerate and etofenprox against local CSFB populations in the Czech Republic in 2015-2018. The LC50 for *lambda*-cyhalothrin in the Prague population in 2015 was 0.0003 μg/cm^2^. Heimbach and Müller [1] recorded in Germany in 2008 the most sensitive population to *lambda*-cyhalothrin with an LC50 of 0.0002 μg/cm^2^ (assessment after 5 h). Compared to the most susceptible CSFB population from Germany, fold resistance of 5 and 20 were calculated in 2018 in the samples from Prague and Potěhy, respectively. However, 100% mortality of CSFB from both populations after application of *lambda*-cyhalothrin in a concentration corresponding to 100% of the recommended field concentration still reflects a satisfactory efficacy of *lambda*-cyhalothrin. The slight decrease in the susceptibility of CSFB to pyrethroids in the Potěhy population might reflect the repeated application of pyrethroids in the last four years in this locality.

The LC50 and LC90 values for pyrethroids in 2015 in the Prague population represent a susceptibility baseline of CSFB prior to the selective pressure of pyrethroids on resistance in the Czech Republic. These baseline data can be used to monitor the susceptibility of CSFB populations to pyrethroids in the Czech Republic. Wide scale monitoring of resistance in 2017 on 7 populations from the Czech Republic (unpublished data) showed high susceptibility to *lambda*-cyhalothrin and *tau*-fluvalinate and slight decrease of susceptibility of one population to *tau*-fluvalinate. In 2018, monitoring of resistance (unpublished data) showed high susceptibility of 10 populations to *lambda*-cyhalothrin and decrease of susceptibility to *tau*-fluvalinate in 13 from the 14 tested populations.

High susceptibility to pyrethroids with very low resistance ratios were also observed in Danish CSFB samples [3]. In contrast to this, evidence for *kdr*-resistance in CSFB was published from Germany [2], where the pyrosequencing diagnostic assay showed a *kdr*-allele frequency of 90-100% in populations expressing high levels of resistance in adult vial tests. Cross-resistance to *lambda*-cyhalothrin, *tau*-fluvalinate, etofenprox and bifenthrin was also shown [2]. In contrast to the German samples, where the survival of CSFB at high concentrations of *lambda*-cyhalothrin correlated well with the presence of the *kdr*-allele, the presence of *kdr*-genotypes in samples from the United Kingdom did not always correlate well with resistant phenotypes. This indicates a possible incidence of an as yet undisclosed, metabolic-based mechanism [3]. Regarding an alarming trend in the spread of CSFB resistance to pyrethroids, including the *kdr*-mutation and metabolic-based mechanisms, it is not recommended to alternate the insecticides on the basis of pyrethroids in anti-resistance strategies. The introduction of pesticides with alternate modes of action is necessary.

### Susceptibility to organophosphates and oxadiazine in the glass vial experiment

Evaluation of the susceptibility of CSFB to insecticides showed a high susceptibility of two local CSFB populations from the Czech Republic to chlorpyrifos and indoxacarb. Up to now, no resistance to chlorpyrifos has been recorded in CSFB. LC50 value 0.01 μg/cm^2^ for chlorpyrifos was established in this paper for the most sensitive sample of CSFB population. Good efficacy of organophosphates was also published against *Brassicogethes* aeneus [8] and *Ceutorhynchus obstrictus* [9]. Chlorpyrifos has recently been recommended for the control of CSFB in the Czech Republic because of the need for alternation with pyrethroid-based insecticides in anti-resistance strategies.

Similarly, no resistance of CSFB or *B. aeneus* to indoxacarb was recorded up to now. LC50 value 0.02 μg/cm^2^ for indoxacarb was established in this paper for the most sensitive sample of CSFB population. Till present, indoxacarb has not been used for control of CSFB in the Czech Republic and is a promising pesticide for anti-resistant strategies to slow down selection of CSFB resistance to pyrethroids. No cross-resistance between indoxacarb and pyrethroids and organophosphates was shown in insecticide-resistant *Spodoptera frugiperda* and *Plutella xylostella* [10]. However, a high resistance level to indoxacarb has been published for *C. obstrictus* from Poland [11].

### Susceptibility to neonicotinoids in the glass vial experiment

This study showed different susceptibilities of both CSFB populations to the neonicotinoids acetamiprid and thiacloprid. Both tested populations of CSFB were highly susceptible to acetamiprid and simultaneously exhibited tolerance to thiacloprid. In the Czech Republic, thiacloprid is not registered for use against CSFB in oildseed rape crops. However, thiacloprid was frequently used in the Czech Republic against *Ceutorhynchus obstrictus* and *Dassineura brassicae* in flowering period of oilseed rape. These applications influence the CSFB larvae escaping the plants after termination of development. Hence, selection pressure of thiacloprid on CSFB larvae in not eliminated.

Extensive surveys have shown that stronger resistance to neonicotinoids has been confirmed in some populations of *Bemisia tabaci* and *Leptinotarsa decemlineata* [12], *Triaulerodes vaporarium* [13], *Nilaparvata* lugens [14] and *Myzus persicae* [15]. A study of *L. decemlineata* resistance to imidacloprid in populations from North America and Europe revealed an approximately 30-fold variation in the LC50 values from ingestion and contact bioassays against neonates [16]. After several years of imidacloprid application, 100-fold [17] and 309-fold [18] resistance ratios for imidacloprid were reported in populations of *L. decemlineata* from Long Island.

No shift to lower susceptibility to thiacloprid has been observed in populations of *B. aeneus* and *C. obstrictus* from several European countries in glass vial experiment with a commercially available thiacloprid formulation, Biscaya OD 240 [19]. According to Zimmer and Nauen [20], the adult vial test with an oil-dispersion-based formulation of thiacloprid resulted in a much better bioavailability compared with technical material.

Resistance to neonicotinoids has been slow to develop compared with other insecticide groups [12]. A target-site resistance has been documented from a selected neonicotinoid resistant laboratory strain of *N. lugens* [21]. However, this mutation has not been found in resistant strains of *N. lugens* collected from the field [22]. From recent evidence, it is clear that neonicotinoid resistance is at present based largely on P450-dependent monooxygenases [23]. It was found that up-regulation of CYP6CM1 in *B. tabaci* confers resistance not only to several neonicotinoids but also to pymetrozine [24]. Puinean et al. [15] published evidence that a P450 gene amplification event is associated with neonicotinoid resistance in *M. persicae*.

No cross-resistance between thiacloprid and acetamiprid or thiamethoxam was recorded for CSFB. Cross-resistance to neonicotinoids was described for several populations and biotypes of the whitefly (*B. tabaci*) [12]. In the present study, the tolerance of CFSB to thiacloprid was detected with preserved susceptibility to acetamiprid and thiamethoxam. Cross-resistance between imidacloprid and thiamethoxam and between acetamiprid and thiamethoxam was observed in *B. tabaci* [25]. Cross-resistance between imidacloprid, thiacloprid and acetamiprid was detected in *N. lugens* [14]. Similarly, cross-resistance between imidacloprid and acetamiprid was detected in *M. persicae* [26].

Little or no cross-resistance of neonicotinoids to older insecticide classes, such as pyrethroids, organophosphates and carbamates, has been observed [27]. One strain of the small brown planthopper (*Laodelphax striatellus*) maintained under laboratory conditions apparently acquired an 18-fold resistance to imidacloprid, without exposure to this insecticide, under strong selection from organophosphates and carbamates, while the unselected strains were fully susceptible to imidacloprid [28].

### Susceptibility to neonicotinoids in the feeding experiment

The feeding experiment showed different susceptibilities of the Prague CSFB population to thiacloprid and thiamethoxam. The samples were highly susceptible to thiamethoxam and tolerant to thiacloprid. The injury to plants in the thiacloprid variant was comparable to the control variant without seed dressing in the feeding experiment. In contrast to this, the seed dressing with thiamethoxam importantly prevented CSFB injury in oilseed rape plants (Table 5). The increase in the average weight of plants in the thiamethoxam variant was observed as a result of the prevention of CSFB injury. A very low susceptibility of CSFB to thiacloprid and also to acetamiprid was reported by Dewar [29].

The experiments proved that seed dressing with thiamethoxam was highly effective for the control of oilseed rape plants in their early growth stages against CSFB. As a consequence of the restriction of seed dressing with three neonicotinoids by the European Commission [5], the injury to oilseed rape plants caused by CSFB increased. It resulted in the increased frequency of the foliar application of insecticides and increased costs for the control of oilseed rape in autumn. The increase in the foliar application of insecticides was also documented in the western part of the EU [6]. Seed dressing with thiacloprid had a very low efficacy due to the CSFB tolerance.

## Conclusions

The study showed good efficacy of the pyrethroids *lambda*-cyhalothrin, deltamethrin, cypermethrin, *tau*-fluvalinate, esfenvalerate and etofenprox, organophosphate chlorpyrifos and oxadiazine indoxacarb against two local CSFB populations in the Czech Republic in 2015-2018. Both the CSFB populations were susceptible to nenonicotinoids acetamiprid and thimethoxam and tolerant to thiacloprid. The CSFB tolerance to thiacloprid was shown in both glass vial and feeding experiments.

## Acknowledgements

We thank Anna Macáková and Jana Vincíková for excellent technical support.

